# Late burst of fork-tail evolution in hirundines (Aves: Hirundininae)

**DOI:** 10.64898/2026.08.17.745338

**Authors:** Masaru Hasegawa

## Abstract

The evolutionary patterns of trait diversification provide insights into the function of the trait. Early burst of trait evolution is often associated with adaptive radiation, rapidly diversifying the trait in response to vacant niches followed by the slowdown of the diversification with niche filling, whereas late burst is more likely to be associated with sexual selection, possibly contributing to reproductive barriers between closely related species. Here, we studied the diversification of tail fork depth through time in hirundines to infer its function, which remains unclear due to the competing two alternative hypotheses: the sexual selection hypothesis, which is a classic explanation of deeply forked tails, proposed that this trait has evolved via sexual selection, which was then challenged by the viability selection hypothesis, which proposed that deeply forked tails have mainly evolved via viability selection for enhancing aerodynamic performance during aerial foraging on large prey. I found a late burst of tail fork depth, but not of bill length, i.e., an index of prey size. The observed pattern is consistent with the sexual selection hypothesis but not with the viability selection hypothesis.

---

The evolutionary patterns of trait diversification through time provide information on the functions of focal traits (Streelman & Danley 2003). An early burst of trait evolution is often found in traits associated with adaptive radiation, in which they evolved rapidly in response to vacant niches followed by the slowdown of the diversification rate with niche filling (e.g., Arbour & López-Fernández 2023). On the contrary, a late burst of trait evolution is more likely to be associated with sexual selection: the diversification of secondary sexual traits can be isolated from ecological diversification, possibly contributing to reproductive barriers between closely related species (e.g., Levoy et al. 2024).

Deeply forked tails of swallows are regarded as a classic example of sexual traits (Møller 1994). Males with experimentally elongated outermost tail feathers (which increases tail fork depth) attract social and extrapair mates, resulting in their reproductive advantage (e.g., Møller 1988; reviewed in Møller 1994; Turner 2006; Romano et al. 2017), explaining the evolution of deeply forked tails. However, an alternative explanation, the viability selection hypothesis, then challenged the sexual selection hypothesis. The viability selection hypothesis proposed that long outermost tail feathers can enhance the aerodynamic performance of swallows and have mainly evolved via viability selection for efficient aerial foraging, perhaps for capturing large, rapid prey (e.g., Norberg 1994; Evans 1998; Buchanan & Evans 2000). A manipulative experiment to test this hypothesis (see Evans & Thomas 1997 for a design) initially seems to support this hypothesis (e.g., Evans 1998), but it is recently shown to be logically flawed because of the lack of consideration for co-evolving compensatory traits (i.e., the manipulation of tail fork depth without taking account of compensatory traits impairs the coordination of deeply forked tails and compensatory traits; thus, it cannot clarify the function of deeply forked tails: reviewed in Hasegawa 2024). Therefore, this manipulative experiment cannot be used to distinguish the two hypotheses (Hasegawa 2024).

A series of macroevolutionary studies, instead, were conducted to infer causes and consequences of the evolution of deeply forked tails of hirundines (e.g., Hasegawa & Arai 2018, 2020a,b, 2022a,b). For example, Hasegawa & Arai (2022a) showed that not foraging habits but opportunities for extrapair mating explains the evolution of forked tails. Hasegawa (2023) showed the coevolution of forked tails and song, that is, an acoustic sexual signal, and Hasegawa (2026) showed that the evolution of tail fork depth explains speciation in hirundines, as predicted by the sexual selection theory. These macroevolutionary studies support the sexual selection hypothesis, but the evidences are still inconclusive (Hasegawa 2025). Further evidence is warranted to clarify the functions of swallows’ tails.

Here, I studied the evolutionary patterns of the diversification of tail fork depth through time in hirundines (Aves: Hirundininae) using disparity through time analysis (Harmon et al. 2008). If sexual selection is an important driver of the diversification as suggested by previous macroevolutionary studies (see above), a late burst of tail evolution would be found. Because evolutionary patterns can be affected by lineage-specific constraints (e.g., presence/absence of out-group competitors, geological events), I also studied the evolutionary patterns of the diversification of other traits. In particular, if viability selection via foraging ability causes the pattern found in tail fork depth, the diversification patterns of tail fork depth should be shared with those of foraging traits. Thus, I studied bill length as an index of prey size as well (e.g., Fitzpatrick 1985; Hasegawa et al. 2016).

## Methods

### Data collection

As before (e.g., Hasegawa 2023, 2025, 2026), I collected morphological data, i.e., body mass (as a measure of body size), wing length (as a measure of wing as a flight apparatus), bill length (as an index of prey size), relative tail fork depth, which are obtained from the monograph of hirundines (Turner & Rose 1994) except for body mass (see below). The main focus here, relative tail fork depth, is calcurated as (male fork depth)/(male central tail feather length) as in previous studies (e.g., Hasegawa & Arai 2017, 2018; Hasegawa 2025). Likewise, I also calcurated female relative tail fork depth (to examine whether female tail fork depth as well as male tail fork depth show late burst of trait evolution). Because bill length as an index of prey size (and male wing length as a measure of flight apparatus) would increase with body size, I focused on residual bill and wing lengths after statistically controlling for body mass as a measure of body size (see the next section). As before (Hasegawa & Arai 2020a), information on body mass was obtained from Turner & Rose (1994) and Dunning (2008).

### Phylogenetic analyses

To evaluate how disparity (i.e., phenotypic diversity) of tail length has changed through time, I conducted disparity through time analysis using the function “dtt” in the R package “geiger” (ver. 2.0.11; Harmon et al. 2008) in R ver. 4.4.2 (R Core Team 2024). Morphological Disparity Index (MDI) was calculated to quantify the overall difference in the observed trait disparity compared with that under Brownian motion by simulating the evolution of tail fork depth 1000 times per tree. Negative MDI values indicate that most disparity originated early in the history of the group, whereas positive MDI values indicate that most disparity originated more recently compared with random walks. To account for phylogenetic uncertainty, a multimodel inference was applied using 1000 alternative trees for hirundines from birdtree.org (e.g., Garamszegi & Mundry 2014; Rubolini et al. 2015; Hasegawa & Arai 2020a,b). Using 1000 alternative trees, the models were fitted to each tree, and the mean MDI and the probability that the observed disparity through time is above/below that of simulated values were estimated (i.e., a two-tailed test; twice the probability of simulations that are above the observed disparity through time out of 1000 simulations × 1000 trees). The disparity of wing length and bill length was also estimated with and without statistically controlling for body mass^(1/3). It should be noted that cube root is used here to match dimensions (i.e., body mass linearly increases with volume, which is expressed in cubic millimeters, mm^3^). In addition, for illustrative purpose, I also computed the phenogram of relative tail fork depth, wing length, and bill length using the function “phenogram” in the R package “phytools” (Revell 2012) using one of the 1000 phylogenetic trees. To depict the evolutionary rates of the phenotypic evolution of each trait through time, Bayesian Analysis of Macroevolutionary Mixtures (BAMM v.2.5.0, Rabosky et al., 2014) was applied using the same phylogenetic tree that was used to depict phenograms (note that each variable was standardized to zero mean and unit variance before analysis to compare the results across traits).

## Results

Disparity through time analysis showed that the disparity of tail fork depth was significantly larger than that expected under Brownian motion (mean MDI = 0.31, *P* = 0.035), indicating a late burst of trait evolution (Figs. 1 & 2 upper panels). When female tail fork depth was used instead of male tail fork depth, the disparity of tail fork depth was not significantly different from that expected under Brownian motion (mean MDI = 0.17, *P* = 0.28).

**Figure 1.**
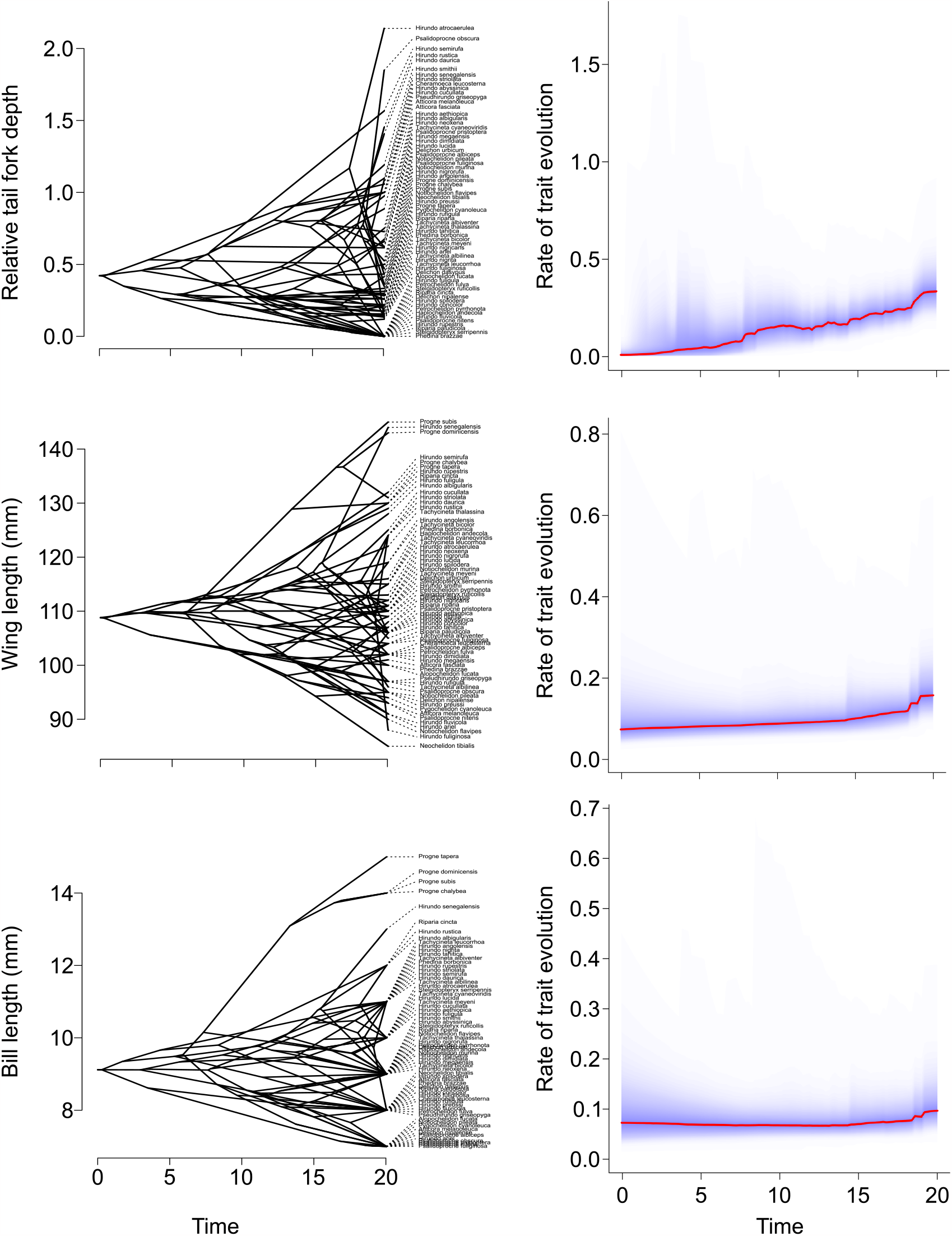
Examples of phenograms (left panels) and corresponding estimated evolutionary rates (right panels) of relative tail fork depth (upper panels), wing length (middle panels), and bill length (lower panels) in hirundines. Phenotypic diversification is rapidly increased near tips in relative tail fork depth (note that rate of trait evolution is standardized to zero mean and unit variance).

**Figure 2.**
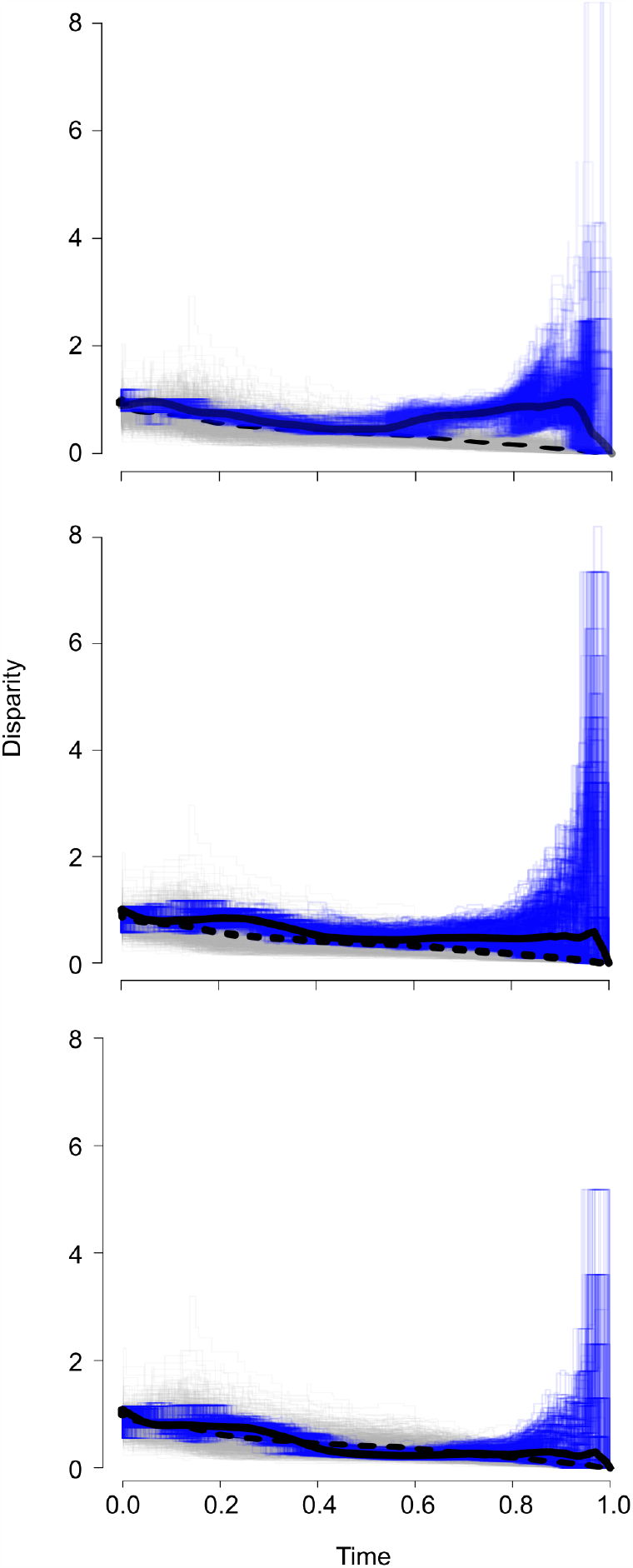
Disparity through time plot for relative tail fork depth (upper panel), wing length (middle panel), and bill length (lower panel) based on 1000 phylogenetic trees used in the current study. The solid lines indicate the observed mean disparity, and the dashed lines indicate the mean simulated disparity under a Brownian model. The thin blue and gray lines indicate individual disparity plots for actual data and simulated Brownian models, respectively. Relative time 0.0 and 1.0 represent the root and the tip of the phylogeny, respectively. The shaded areas on each plot indicate the 95% confidence interval for the simulations. See text for the formal statistics.

The disparity of wing length did not show a significant difference from that expected under Brownian motion (i.e., not significantly different from zero; MDI = 0.22, *P* = 0.17; Figs. 1 & 2 middle panels). When residual wing length was used to statistically control for a measure of body size, body mass (which also did not show a significant difference from that expected under Brownian motion: MDI = 0.11, *P* = 0.51), showed a similar nonsignificant pattern (MDI = 0.28, *P* = 0.06). Bill length, which is an index of prey size (see Methods), also did not show a significant difference from that expected under Brownian motion (MDI = 0.04, *P* = 0.85; Figs. 1 & 2 lower panels). This was also the case when residual bill length was used to statistically control for body mass (MDI = 0.22, *P* = 0.10).

## Discussion

The main finding of the current study is that the evolutionary patterns of the diversification of male tail fork depth in swallows showed a late burst of trait evolution. This finding is contrary to the pattern found in bill length, that is, an index of prey size, which did not differ from that expected from Brownian motion (i.e., random walks). A nonsignificant trend of wing length is predictable, because a long wing is a compensatory trait for deeply forked tails (e.g., Møller et al. 1995; Møller 1996; Hasegawa 2024), and thus the pattern of wing length would show a trend similar to that of tail fork depth, particularly after correcting for body size (i.e., after excluding variance explained by body mass, which is irrelevant to a late burst). Therefore, the obtained pattern here supports the sexual selection hypothesis rather than the viability selection hypothesis. Sexual selection on tail fork depth would be particularly important at later stages of lineage diversification, possibly via its role in the formation of reproductive barriers and reinforcement (e.g., Levoy et al. 2024; reviewed in Streelman & Danley 2003).

A series of macroevolutionary studies supports the sexual function of forked tails; 1) measures of foraging costs linearly increase with fork tail depth (Hasegawa et al. 2016; Hasegawa & Arai 2017; Hasegawa 2025); 2) forked tails coevolved with song, that is, an acoustic sexual signal (Hasegawa 2023); 3) the perceived, rather than physical, size of forked tails matters (Hasegawa & Arai 2020a); 4) the indices of sexual selection, not those of foraging habit, explain the evolution of deeply forked tails (Hasegawa & Arai 2020b, 2022a); 5) extinction risk increased with tail fork depth in our changing world (Hasegawa & Arai 2021, 2022b). Finally, a recent study of trait-dependent lineage diversification has shown the association between the diversification of tail fork depth and speciation as predicted by sexual selection theory (Hasegawa 2026). The current study of time-dependent trait diversification is consistent with previous studies focusing on different aspects of sexual selection theory, reinforcing the idea that sexual selection explains the evolution and diversification of swallows’ tails.

A caveat of the current study is the correlational nature of disparity through time analysis. The confounding effects of unmeasured correlates of tail fork depth cannot be completely ruled out. However, even in this case, it should be noted that a late burst of bill evolution was undetected, suggesting that a late burst of tail evolution was not due to the diversification of prey size. No detectable late burst of fork-tail evolution in female swallows, which should be less affected by sexual selection, reinforces this perspective. Therefore, the negative evolutionary relationships between prey size (or its correlate, bill length) and tail fork depth found in this study system (e.g., Hasegawa et al. 2016; Hasegawa 2025) would not imply the diversification of forked tails adapted to differential foraging niches but rather suggest the foraging cost of deeply forked tails, which prevents hirundines that forage on large prey from having deeply forked tails or vice versa, as suggested before (e.g., extinction risk increases with tail fork depth in our changing world of “insect apocalypse”; Hasegawa & Arai 2021). Some manipulative experiments support this explanation (e.g., Cuervo & Møller 2006), but further evidence is needed (because of the lack of consideration for compensatory traits: Hasegawa 2024; see Introduction). Likewise, the direct causation behind a late burst of male tail-fork evolution remains to be clarified, which might be accomplished by using genomic analyses across hirundines (e.g., see Shields et al. 2024 for the role of deeply forked tails in the within-species reproductive isolation in the barn swallow).

In summary, a late burst of evolution of male tail fork depth (but not that of bill length) was found here, which is consistent with the sexual selection hypothesis but not with the viability selection hypothesis. Thus, although confounding factors cannot be ruled out in this correlational study, the observed patterns provide an additional support for the importance of sexual selection on the evolution and diversification of swallows’ tails (to those found in previous macroevolutionary studies; see above). Disparity through time analysis could be affected by several factors, including evolutionary events of outgroups (e.g., emergence and diversification of swifts, i.e., another hyper-aerial insectivores, in the current case) and geological events (e.g., geographic isolation of suitable habitats), but the diversification pattern of the focal trait (i.e., tail fork depth, here) together with other functional traits related to the focal trait (i.e., bill size as an index of prey size, here) could provide better insights into the function of the focal trait.

## Acknowledgments

I thank Dr Emi Arai for valuable comments and thank Dr Shumpei Kitamura and his lab members at Ishikawa Prefectural University for their kindest advices. I am grateful to Dr Angela Turner for her kindly support on the valuable information on swallows. I was supported by KAKENHI grant (JSPS, 22J40066).

## Conflict of interest

I have no competing interests.

---

**Table S1.**
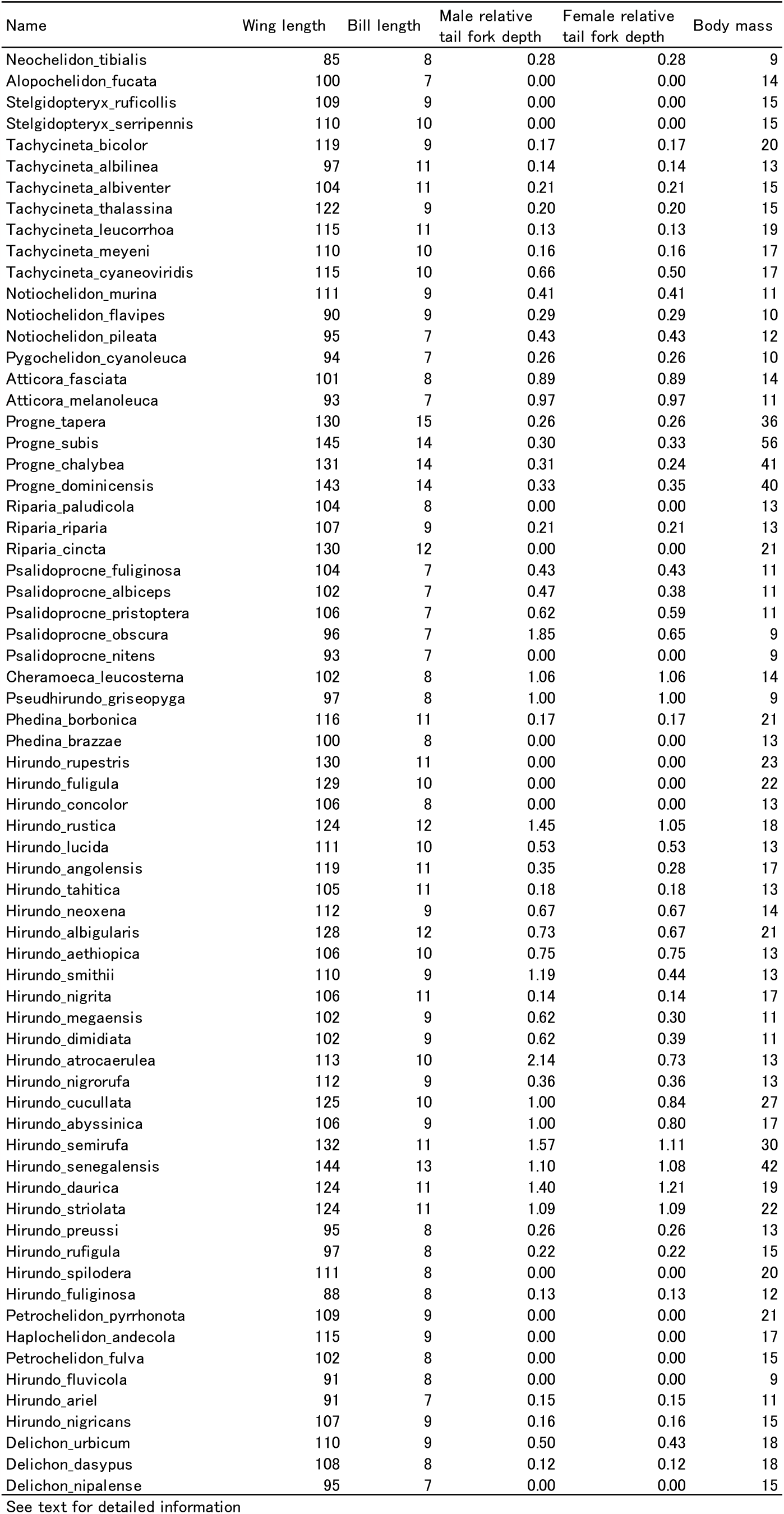
Dataset of the current study (*n* = 68).

## Notes

### Competing Interest Statement

The authors have declared no competing interest.

